# A novel microporous biomaterial vaccine platform for long-lasting antibody mediated immunity against viral infection

**DOI:** 10.1101/2024.01.30.578038

**Authors:** Daniel P. Mayer, Mariah E. Neslon, Daria Andriyanova, Olivia Q. Antao, Jennifer S. Chen, Philip O. Scumpia, Westbrook M. Weaver, Stephanie Deshayes, Jason S. Weinstein

**Author notes:** Department of Internal Medicine, Lankenau Medical Center, Wynnewood, PA 19096. Co-corresponding authors Lead Contact: Jason S. Weinstein Cancer Center, G-1216 Rutgers New Jersey Medical School 205 South Orange Avenue, Newark, NJ 07103.

## Abstract

Current antigen delivery platforms, such as alum and nanoparticles, are not readily tunable, thus may not generate optimal adaptive immune responses. We created an antigen delivery platform by loading lyophilized Microporous Annealed Particle (MAP) with aqueous solution containing target antigens. Upon administration of antigen loaded MAP (VaxMAP), the biomaterial reconstitution forms an instant antigen-loaded porous scaffold area with a sustained release profile to maximize humoral immunity. VaxMAP induced CD4+ T follicular helper (Tfh) cells and germinal center (GC) B cell responses in the lymph nodes similar to Alum. VaxMAP loaded with SARS-CoV-2 spike protein improved the magnitude and duration of anti-receptor binding domain antibodies compared to Alum and mRNA-vaccinated mice. A single injection of Influenza specific HA1-loaded-VaxMAP enhanced neutralizing antibodies and elicited greater protection against influenza virus challenge than HA1-loaded-Alum. Thus, VaxMAP is a platform that can be used to promote adaptive immune cell responses to generate more robust neutralizing antibodies, and better protection upon pathogen challenge.

## 1. INTRODUCTION

High-affinity neutralizing antibodies and long-lived memory B cells generated in a germinal center (GC) reaction are hallmarks of the humoral response (1, 2). Long-lived plasma cells generated during a primary infection or vaccination continually secrete antibodies conferring the first wave of protection against re-infection (3). When these antibodies fail to control the re-infection, pathogen-experienced memory B cells differentiate into short-lived antibody secreting cells or secondary germinal centers to expedite the recall response. The efficacy of long-lived plasma cells and memory B cells to confer protection upon subsequent challenges depend on their development during the initial immune insult (4). Coordination of immune cell types and cytokines during the primary immune response, as well as sustained release of antigens is required for the generation of protective responses (3, 5, 6). Generating long-lived, neutralizing antibody responses against viral entry proteins, such as the receptor binding domain (RBD) of the spike protein of SARS-CoV-2 or hemagglutinin (HA1) of influenza virus can protect people from disease. Most current vaccines, including COVID-19 vaccines, induce transient antigen expression, which may not provide necessary immune response for driving robust recall responses (6, 7). Thus, there is a strong need for new vaccine strategies or platforms to provide sustained antigen release.

The development of a vaccine delivery system that allows for control of antigen release kinetics should allow for an improved pathogen-specific recall response. It has been shown that the magnitude, functionality, and phenotype of antigen-specific CD4+ and CD8+ T-cell responses can be shaped by controlled release of antigen and immunomodulatory biologics over several weeks and provide stronger immunization compared to bolus injections (8). Due to their tunability, bioengineered vaccines with sustained antigen release may allow for improvements in tailoring the immune response against a particular pathogen. Bioengineered immunogens have been developed to elicit improved vaccine responses, by sustaining immune responses longer than typically possible with standard adjuvants. Using hydrogel scaffolds made of synthetic and natural materials as vaccine delivery platforms represent another way to sustain release of antigen for vaccine delivery systems to modulate an immune response (8–16). The physical properties of the material used in developing vaccine delivery platforms are critical for controlling and regulating the activity of the immune response (17–20). Different materials can alter antigen uptake, immune cell migration to draining lymph nodes, and immune cell activation. Polymeric microspheres and aluminum hydroxide, for example, have been widely investigated as vaccine carriers to provide sustained release (21–23). Recently, a new class of biomaterial scaffolds – granular hydrogels - were created with a hyper-porous geometry to allow for optimal cell infiltration while minimizing the host foreign body response (24). Microporous annealed particle (MAP) hydrogels are a flowable suspension of concentrated viscoelastic microspheres ‘building blocks’ that transition *in situ* to form a robust porous scaffold (23, 25–27). Significantly more immune cells can be exposed to the loaded antigen as they migrate through the hyper-porous space around annealed MAP building blocks, and as the scaffold containing the antigen slowly degrades, antigen will be released from the scaffold.

Early formulation of MAPs have shown the ability to elicit tissue regeneration by activating antigen-specific antibody responses in mice loaded with antigen (25). Proof of concept work showed that MAP loaded with immunogens can elicit antigen specific immunity and tuning hydrogel stiffness can serve an adjuvant role, enhancing antibody responses (27). However, given the need for microfluidic fabrication, with the necessity to incorporate antigenic material during manufacturing, the ability to develop the MAP as a tunable immunization platform for medical purposes was limited. Herein, we leveraged the unique properties of the MAP platform to develop a next generation vaccine delivery system (VaxMAP) with improved adaptive immune responses without the need for microfluidic fabrication. Various immunogens were easily loaded into lyophilized MAP particles post-fabrication with complete control over loading capacity and no immunogen loss providing a vaccine delivery scaffold with a two-stage release profile (**Fig. 1**). This two-stage release profile is characterized by an initial burst release for antigen exposure to prime immune responses followed by a slower and sustained release for long-term memory antibody and cellular responses. We demonstrated the versatility of the VaxMAP platform to immunize mice against two key human viral pathogens: SARS-CoV-2 and influenza. We explored VaxMAP’s ability to induce a prolonged antibody production targeting different viruses: SARS-CoV-2 (spike) and influenza (HA1). We showed that immunization with spike-loaded VaxMAP greatly enhanced the magnitude and duration of anti-RBD antibodies compared to spike loaded in Alum. Similarly, VaxMAP loaded with HA1 elicited better antibody responses against hemagglutinin and protection against influenza virus infection than Alum. Thus, VaxMAP represents an injectable biomaterial platform to enhance humoral immunity for traditional vaccines.

**Fig. 1.**
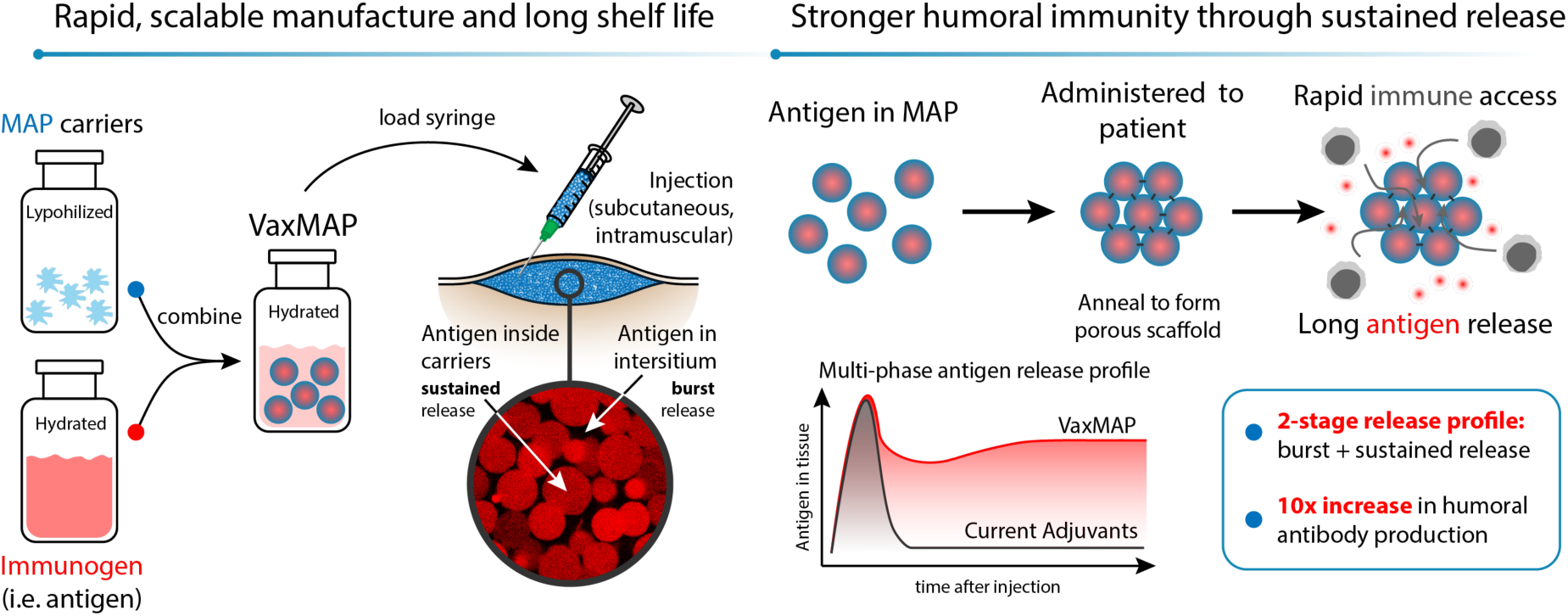
VaxMAP scaffold provides a rapid way to load an immunogen at the desired concentration with a two-stage release profile. The hydrogel microspheres that constitute MAP are provided lyophilized for longer shelf-life stability and are reconstituted with an immunogen at time of use. While the hydrogel microspheres are swelling, part of the antigen is passively diffusing within the hydrogel microspheres, whereas some of the antigen is remaining within the interstitial space between particles. VaxMAP has pores that are large enough for the immune cells to immediately migrate into the scaffold prior to its degradation. Its double-stage release profile provides an initial burst release for antigen exposure to prime immune responses followed by a slower and sustained release for long-term memory antibody and cellular responses.

## 2. MATERIALS and METHODS

### 2.1. Hydrogel microsphere synthesis

To enable hydrogel microsphere formation, two aqueous solutions were prepared. One solution contained a 10% w/v 4-arm PEG-vinyl sulfone (PEG-VS, 20 kDa, NOF America) in 300 mM triethanolamine (Spectrum Chemical), pH 7.8, pre- functionalized with 500 μM K-peptide (Ac-FKGGERCG-NH_2_) (Bachem), 500 μM Q-peptide (Ac-NQEQVSPLGGERCG-NH_2_) (Bachem) and 1 mM RGD (Ac-RGDSPGERCG-NH_2_) (Bachem). The other solution contained a 7.5 mM di-cysteine modified Matrix Metalloprotease (MMP) peptide substrate (Ac-GCRDGPQGIWGQDRCG-NH_2_) (Bachem). RGD, K- and Q-peptides were added for cell adhesion and secondary annealing as previously reported (23, 25–28). The pre-prepared aqueous PEG-VS and MMP-peptide solutions were mixed in a 1:1 volume ratio, forming the “pre-hydrogel solution”. The pre-hydrogel solution was then immediately injected in mineral oil (with 1% wt Span80) and stirred for 2 hours at 1250 rpm at room temperature. Within the droplets of pre-hydrogel solution, the polymer (PEG-VS) and the peptides covalently bind to one another through a Michael addition reaction between the vinyl sulfone (VS) and the thiol groups. As the reaction progresses, the liquid droplets are converted to hydrogel microspheres. The formulation was designed with a specific molar ratio between the VS groups from the PEG-VS and the thiols from the synthetic peptides such that excess VS remained because of its role in the chemical-annealing reaction. The particles were left to settle down overnight before decanting oil. The particles were then purified by tangential flow filtration (TFF) using 95% isopropyl alcohol (IPA) and lyophilized.

### 2.2. Particle size analysis

The size of the hydrogel microspheres was determined using a laser diffraction particle size analyzer (LS 13320 Beckman-Coulter). Briefly, lyophilized particles were reconstituted at a concentration of 20 mg/mL in phosphate buffered saline (PBS, 10 mM, pH 7.4). The particles were left to swell for 24 hours at room temperature before testing. Then, the particles were diluted 1:10 in water and added to a Beckman LS 13320 laser diffraction particle size analyzer. A volume-weighted distribution curve was obtained from which the mean, mode, and standard deviation were calculated. The LS 13320 machine was verified before use using Garnet G15 control particles provided by the manufacturer (Beckman-Coulter).

### 2.3. Spike antigen loading

Lyophilized hydrogel microspheres were resuspended at a concentration of 20 mg/mL in phosphate buffered saline (PBS, 10 mM, pH 7.4) containing 1.25 mg/mL of SARS-CoV-2 spike protein (SARS-CoV-2 spike S1+S2 ECD (R683A, R685A, F817P, A892P, A899P, A942P, K986P, V987P)-His recombinant protein, Sinobiological, cat # 40589-V08H4). The particles were left to swell for 24 hours at 2-8°C before use. At this point, the volume fraction (VF) of the particles was 100% (1 mL/mL). 0.8 mL of spike-loaded hydrogel microspheres were then loaded in 1-cc sterile syringe.

### 2.4. Hemagglutinin antigen loading

1.2 mg of influenza hemagglutinin (HA) protein (Influenza A H1N1 (A/Puerto Rico/8/1934) hemagglutinin protein, Sinobiological, cat # 11684-V08H) were dissolved in phosphate buffered saline (PBS, 10 mM, pH 7.4), and then washed to remove glycerol and concentrated down to 1.25 mg/mL using PES concentrator tubes (MWCO 10 kDa) by centrifugation at 4°C at 8000 rcf for 2 hours. Lyophilized hydrogel microspheres were resuspended at a concentration of 20 mg/mL in the PBS solution (pH 7.4) containing 1.25 mg/mL of HA protein. The particles were left to swell for 24 hours at 2-8°C before use. At this point, the volume fraction (VF) of the particles was 100% (1 mL/mL). 0.8 mL of HA-loaded hydrogel microspheres were then loaded in 1-cc sterile syringe.

### 2.5. Chemical annealing

Immediately prior to use (e.g., for testing or injection in animals), the hydrogel microspheres (with or without antigen) were annealed with a PEG-dithiol crosslinker (3.4 kDa, NOF America). Briefly, a solution of 1 mM PEG-dithiol was prepared in phosphate buffered saline (PBS, 10 mM, pH 7.4). Then, 200 µL of the 1 mM crosslinker solution was pulled up into an empty syringe and mixed thoroughly with 800 µL of hydrogel microspheres to bring the volume fraction down to 80%, the antigen concentration down to 1 mg/mL (when loaded with an antigen), and the crosslinker concentration down to 0.2 mM. The final annealed product is referred to as MAP when no antigen was loaded, as VaxMAP(S) when spike was loaded, and as VaxMAP(HA) when hemagglutinin was loaded.

### 2.6. Mechanical testing

The elastic modulus before and after annealing of the hydrogel microspheres (no antigen) was measured in a compressive test using a 3342 Instron. Briefly, lyophilized hydrogel microspheres were reconstituted at a concentration of 20 mg/mL in phosphate buffered saline (PBS, 10 mM, pH 7.4). The hydrogel microspheres were left to swell for 24 hours at room temperature before testing. Immediately after mixing the particles with the annealing crosslinker (PEG-dithiol) as described above, 80 µL of the particles being annealed were pipetted in a mold made of silicon spacer placed onto a glass slide. A second glass slide was placed on top of the spacer to protect the sample from evaporation. One hour after annealing, mechanical testing was performed using a 3342 Instron instrument with a 2.5-N load cell. A 3-mm diameter anvil descended into the sample at 5 mm/min. The test was completed when the force reached 0.2 N. Compressive force and extension were measured. The raw force/displacement data from the Instron were converted to stress/strain curves to calculate the compressive elastic modulus (EM). For the non-annealed condition, the particles were mixed with 200 µL of PBS solution that did not contain the PEG-dithiol crosslinker and served as a control. Four replicates per condition (N = 4) were performed.

### 2.7. Conjugation of a fluorescent dye to Immunoglobulin G (IgG)

Human Immunoglobulin G (IgG) (Millipore-Sigma) was fluorescently labeled with CF^TM^ 568 maleimide dye (CF568) (Millipore-Sigma). Briefly, the protein (25 mg) was dissolved at a concentration of 67 µM in 2.5 mL of phosphate buffered saline (PBS, 10 mM, pH 7.4). Then, 180 µL of a 10 mM tris(2-carboxyethyl)phosphine hydrochloride (TCEP) (Millipore-Sigma) solution was added to IgG solution to reduce disulfide bonds. After 20 min, 114 µL of CF568 dye (10 mM in DMSO) was added to the mixture. The solution was stirred for 2 hours at room temperature protected from light. The protein was purified with water using a 10 mL 7 kDa MWCO Zeba spin desalting column (Thermo Fisher Scientific), and then by dialysis using a 3.5 kDa MWCO regenerated cellulose Spectra/Por®3 dialysis membrane (Spectrum Labs). The protein (CF568-labeled IgG) was recovered by lyophilization (24 mg).

### 2.8. Protein release

CF568-labeled IgG (1.25 mg/mL) was loaded to MAP following the same procedure described above for the antigens (spike and HA). MAP(IgG) was annealed immediately prior to the release experiment following the procedure described above for chemical annealing. Then, 300 µL of annealed MAP(IgG) was placed in a well insert in a 24-well plate, and 600 µL of PBS (10 mM, pH 7.4) were added to the well. The plate was incubated at 37°C for 25 days. At specific time intervals during the release, the solution was collected from each well and replaced with fresh PBS. After 21 days of release, trypsin at a concentration of 0.75 µg/mL was added to the well to degrade MAP and induce the release of the remaining loaded protein. The concentration of released protein was measured by fluorescence spectroscopy (excitation 579 nm, emission 603 nm) using a calibration curve from known concentrations of CF568-labeled IgG.

### 2.9. Animals

Mice were housed in pathogen-free conditions at the Rutgers New Jersey Medical School. C57BL/6J (B6) were purchased from NCI managed colony at Charles River. Male and female mice were used in experiments with sex-matched controls. All animals were used starting at 8 weeks of age, with approval for all procedures given by the Institutional Animal Care and Use Committee of Rutgers New Jersey Medical School.

### 2.10. Murine immunization

Mice were injected subcutaneously on the right flank by the base of the tail with recombinant antigen. For recombinant SARS-CoV-2 spike experiments, mice received 50 µg spike in either i) VaxMAP(S) (50 µL, immediately after annealing), ii) Alum (100 µL, Thermo Fisher Scientific) or iii) PBS (100 µL). For recombinant Influenza, mice received 50 µg HA1 (Sino Biologics) in either i) VaxMAP(S) (50 µL, immediately after annealing), ii) Alum (100 µL, Thermo Fisher Scientific) or iii) PBS (100 µL). Animals were sacrificed at indicated time points post immunization (p.i.) and harvested organs were processed for flow cytometry. To evaluate the antigen-recall response, a cohort of immunized mice were re-challenged 65 days after the first immunization with 100 µl s.c. injection on the left flank with Alum (mixed with 50 µg spike protein). These mice were euthanized 7 days after the re-challenge.

### 2.11. Influenza infections

Mice were anesthetized with 87.5mg/kg Ketamine/ 12.5 mg/kg Xylazine cocktail in 100 µL of PBS prior to infection. 60 days after immunization, mice were infected intranasally with 30 µL 2×10^6^ PFU of influenza A/Puerto Rico/8/1934 H1N1 (PR8) in PBS using a p200 pipette as determined by TCID_50_ (Rodriguez, Nogales, and Martinez-Sobrido 2017). The survival rate and weight loss and recovery were monitored for 14 days post-infection.

### 2.12. Flow cytometry

Tissues were homogenized by crushing with the head of a 1-ml syringe in a Petri dish followed by straining through a 40-µm nylon filter. Ammonium–chloride–potassium buffer was used for red blood cell lysis. To determine cell counts, count beads were used at a concentration of 10,000 beads/10 µL and added directly to sample following staining. Cell count is normalized by dividing input bead count by cytometer bead count and multiplied by the dilution factor. Antibodies used for flow cytometry staining are listed in **Table S1**. Staining was performed at room temperature (25°C) with 30 min of incubation. Stained and rinsed cells were analyzed using a multilaser cytometer (LSRII, or Fortessa; BD Biosciences or Attune; Invitrogen).

### 2.13. Enzyme-linked immunosorbent assay (ELISA)

2 µg/mL of spike RBD antigen (SARS-CoV-2 spike RBD-His recombinant protein, Sinobiological, Cat # 40592-V08H) or HA antigen was coated onto 96-well ELISA plates overnight at 4°C. Plates were then washed with PBST (PBS + 0.1% Tween 20) and blocked with 3% milk solution for 1 hour at room temperature. Serum was serially diluted in the plates starting at a 1:100 dilution and incubated for 2 hours followed by washes with PBST. Normal mouse serum (obtained from SouthernBiotech, cat # 0050-01) was also used as a negative control. Anti-mouse secondary antibody (goat anti-mouse IgG, human ads-HRP, SouthernBiotech, cat # 1030-05) at 1:8000 dilution in blocking buffer was incubated at room temperature for 1 hour. The plates were washed, and SigmaFast OPD (Millipore) solution was added. After 10 minutes, 3 M HCl solution was added to stop the reaction, and OD at 490 nm was measured on a microplate reader. The binding response (OD_490_) was plotted against the dilution factor in log10 scale as the dilution-dependent response curve. The reciprocal binding antibody titers were calculated based on the area under curve of the dilution-dependent response to quantify the potency of the serum antibody binding to spike RBD or HA antigen.

Serum samples from mice immunized with a single dose of mRNA-1273 vaccine (1 µg) were previously performed by Dr. Eisenbarth and collaborators (30). Briefly, used vials of mRNA-1273 vaccine were obtained from Yale Health within 6 hours of opening. All vials contained less than one full dose per vial, and no vaccines were diverted for the purpose of this study.

### 2.14. Enzyme-linked immunosorbent spot (ELISpot)

SARS-CoV-2 spike antigen was coated at a concentration of 2 μg/mL in carbonate buffer on 96-well MultiScreen HTS IP Filter plates (Millipore) overnight at 4°C. Plates were blocked with complete RPMI (10% heat-inactivated FBS, 1% Penicillin/Streptomycin, 2 mM L-glutamine, 1 mM sodium pyruvate, 10 mM HEPES, 55 μM 2-mercaptoethanol) for 2 hours at 37°C. Bone marrow cells were isolated from the left femur + tibia of mice. Red blood cells were lysed with RBC Lysis Buffer for 2 minutes. Cells were resuspended in complete RPMI and plated in duplicate at three dilutions (1/5, 1/10, and 1/20 of total bone marrow cells) for 20 hours at 37°C. Plates were washed six times with PBST (0.01% Tween-20), followed by incubation with anti-mouse IgG-Alkaline Phosphatase (SouthernBiotech, 1030-04) in PBS with 0.5% BSA for 2 hours at room temperature. Plates were then washed three times with PBST and three times with PBS. Spots were developed with Vector® Blue Substrate Kit (Vector Laboratories, SK-5300) and imaged and counted with an C6ImmunoSpot analyzer (Cellular Technology Limited).

### 2.15. Hemagglutinin inhibition assay

250 µl of Turkey red blood cells were diluted in 10mL 1X PBS. 50 µl of diluted RBCs was then added to experimental wells on a 96-well V bottom plate. Influenza-A was prepared using a two-fold serial dilution and then different dilutions were added to specific wells. Sera samples from mice immunized with Alum-(HA) or VaxMAP(HA) were then added to the wells containing virus and incubated at room temperature for 30 minutes to assess the presence of hemagglutination.

### 2.16. Statistical analysis

All statistical analysis was performed using GraphPad Prism 10 software. Specifically, two-tailed t-test were used to determine statistical significance, assuming equal sample variance for each experimental group when comparing individual groups. P < 0.05 was considered statistically significant.

## 3. RESULTS and DISCUSSION

### 3.1. MAP scaffold fabrication and characterization

Microporous Annealed particles (MAP) are made of hydrogel microspheres that are comprised of 4-arm-poly(ethylene glycol) (PEG)-vinyl sulfone (PEG-VS) backbone crosslinked with a synthetic peptide that imparts biodegradability via protease sensitivity to matrix metalloproteinases (MMPs) (**Figure 2A**). The formulation was designed such that an excess of vinyl sulfone (VS) groups remained after hydrogel microsphere synthesis to enable *in situ* annealing to form a structurally stable scaffold with cell-scale microporosity. The manufacturing process for MAP production has been proven to be rapid and highly scalable. Furthermore, MAP is provided lyophilized which allows for longer shelf-life stability and can eventually facilitate deployment in traditional and austere environments. Given the fact that the deployment of current vaccine technologies, including mRNA vaccines, has been riddled with various challenges, such as severe supply chain constraints, that limit worldwide vaccination efforts, the development of a shelf stable and stockpiled vaccine is in dire need.

**Fig. 2.**
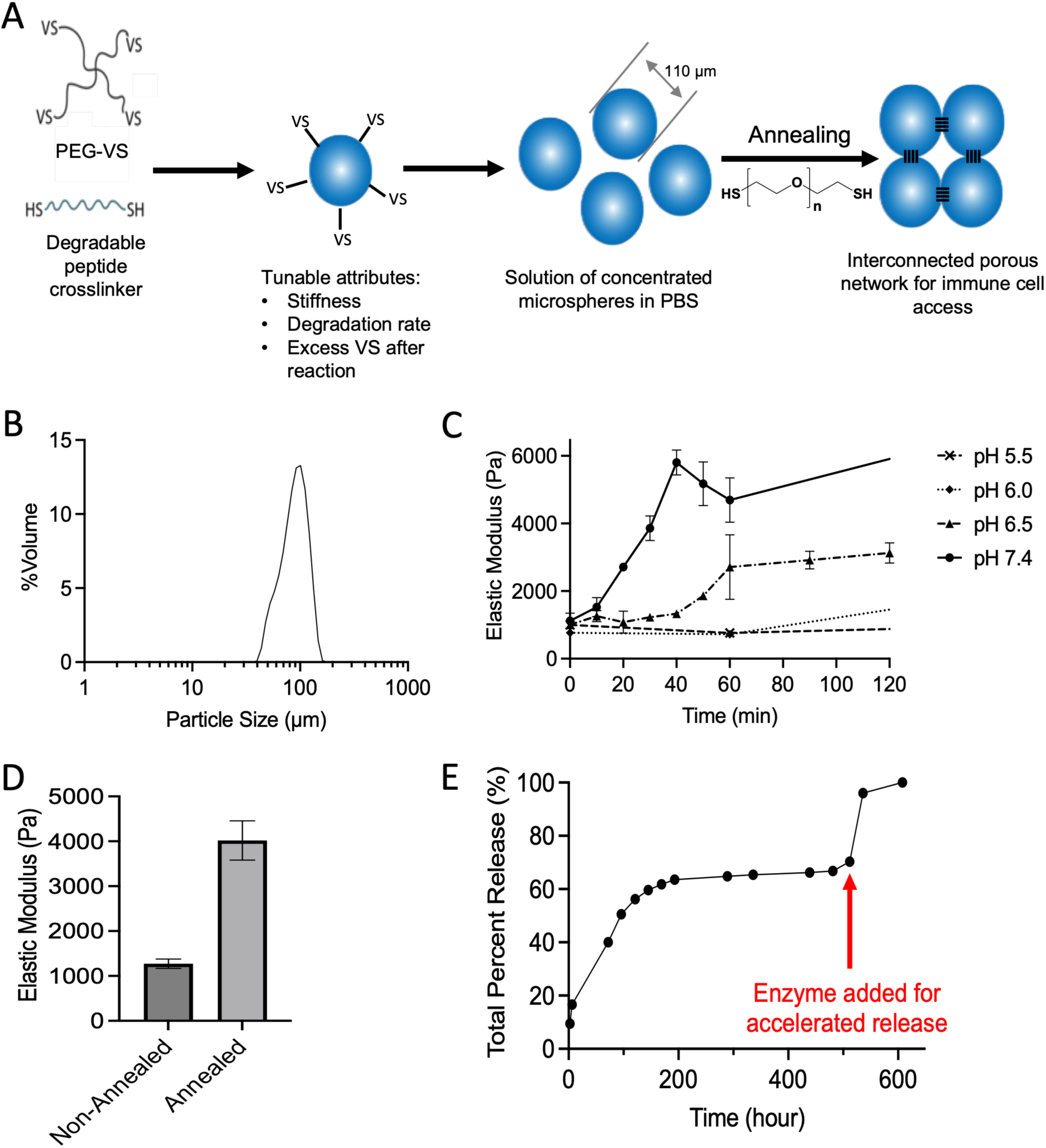
**(A)** MAP hydrogel microsphere fabrication and chemical annealing. The hydrogel microspheres are obtained by a crosslinking reaction between the vinyl sulfone (VS) groups on the PEG polymer (PEG-VS) and the thiol (SH) groups on the degradable peptide through Michael addition in a water-in-oil emulsion. Immediately prior to use, MAP is mixed with a PEG-dithiol crosslinker leading the excess VS groups on neighboring microspheres to react with the thiol groups (chemical annealing) to form an interconnected porous network. **(B)** Particle size distribution (volume weighted) after lyophilization, reconstitution and swelling of the hydrogel microspheres in PBS (pH 7.4). **(C)** Annealing kinetic of MAP at different pHs. Increasing pH accelerates the annealing reaction. **(D)** Compressive elastic modulus before and after annealing the hydrogel microspheres using PEG-dithiol (final concentration: 0.2 mM) at pH 7.4 and at a volume fraction of 80%. Mean ± SD (N = 4). **(E)** Release profile of a surrogate protein from MAP. The red arrow indicates the addition of trypsin to accelerate cargo release.

After lyophilization, reconstitution and swelling in an aqueous phosphate buffered saline (PBS) solution (pH 7.4), the hydrogel microspheres displayed a mean size of 110 µm, a mode size of 105.9 µm, a standard deviation of 23.9 µm resulting in a coefficient variation (CV) of 23.5% (**Fig. 2B**). The polydispersity index (PDI) was then calculated to be 0.055. Particles with PDI values smaller than 0.1 are considered to have a narrow size distribution.

The hydrogel microspheres can be annealed (crosslinked) to each other with a variety of methods to generate a stable porous MAP scaffold. The resultant annealed MAP scaffold displays a functional open pore structure with microscale porosity (pore size: 10 - 50 μm) to allow for free movement of cells and biologics (26). This hyper-porous structure is essential to the scaffold’s inherent lack of a foreign body response that renders MAP non-immunogenic when no antigens are embedded – unlike any other current materials or particle delivery systems (31). Since our previous methods to anneal the hydrogel would not work subcutaneously or would be too expensive for distribution in a vaccine setting, we developed a new chemical annealing method using excess vinyl sulfone groups (VS) and the addition of an annealing agent, PEG-dithiol (3.4 kDa) (**Fig. 2A**). No additional modification of the hydrogel microspheres was required. The thiol groups of PEG-dithiol react with the excess VS groups of the particles to form a thioether bond through a thiol-ene Michael addition. This reaction kinetic is highly dependent upon pH. An increase in pH accelerates the completion of the annealing reaction (**Fig. 2C**). At pH 7.4, it was determined to be complete within an hour. This chemical annealing method offers several advantages: (i) simple, (ii) safe (PEG-dithiol 3.4 kDa is not cytotoxic at least up to 1 mM; data not shown), (iii) uses an inexpensive crosslinker, (iv) an adjustable kinetic and (v) it is not limited by tissue depth as for photo-annealing (white light does not penetrate deep tissues). The compressive elastic modulus of annealed hydrogel microspheres (MAP) was more than 3-fold compared to non-annealed microspheres demonstrating that when the scaffold is annealed, it can better resist the applied force creating a more stabilized scaffold (**Fig. 2D**).

### 3.2. Antigen loading and release from VaxMAP

We engineered MAP as a vaccine delivery platform by adding the spike (S) protein of SARS-CoV-2 within the hydrogel microspheres. The spike protein is a large type I transmembrane protein containing two subunits S1 and S2. S1 contains the receptor binding domain (RBD) which is responsible for recognizing the cell surface receptor and S2 contains elements for membrane fusion. The S antigen was loaded simply by mixing a buffered solution containing the protein at a desired concentration with lyophilized hydrogel microspheres post fabrication. While the particles are swelling, part of the antigen is passively diffusing within the hydrogel microspheres, whereas some of the antigen is remaining within the interstitial space between particles (**Fig. 1**). This loading method offers a very quick way to entrap the antigen with no loss of antigen and complete control over the antigen concentration.

The release profile of a surrogate fluorescently labeled protein with a similar molecular weight to the spike protein has been investigated (**Fig. 2E**). During the first stage of the release, the protein loaded in the interstitial space between particles is rapidly released by passive diffusion from the scaffold (which represents about 60% of the total protein), while during the second stage, the protein entrapped within the particles is slowly released following the degradation rate of the particles. Trypsin was added to purposely accelerate the degradation of the scaffold during the second stage of the release which led to a sudden release of the remaining protein from the particles (**Fig. 2E**). This double-stage release profile provides an initial burst release for antigen exposure to prime immune responses followed by a slower and sustained release for long-term memory antibody and cellular responses.

At time of use, immediately prior to injection, the spike-loaded hydrogel microspheres are chemically annealed using the PEG-dithiol crosslinker to create a stabilized scaffold referred here as VaxMAP(S).

### 3.3. VaxMAP promotes an adaptive immune response

As spike was released from MAP in two phases, we investigated the ability of VaxMAP(S) to induce a primary immune response in a murine immunization model. Mice were immunized subcutaneously (s.c.) with a single dose of 50 µg of spike added to either VaxMAP(S), Alum (Alum(S)) or PBS (PBS(S)). To assess the immune response following immunization, we examined the primary Tfh and GC B cells formation in the draining inguinal lymph nodes (iLN) at days 10, 28 and 60 post immunization (p.i.). There was a similar increase in the frequency of Tfh cells (CD4^+^CD44^+^Ly6c^lo^PSGL-1^lo^CXCR5^+^PD-1^+^) in VaxMAP(S) and Alum(S) immunized mice at days 10 and 28 compared to PBS(S), yet at day 28 Tfh cells were slightly elevated in VaxMAP(S) compared to Alum(S) (**Fig. 3A**). By day 60, the frequency of Tfh cells in both groups was reduced to that of the PBS(S) treated mice. Likewise, the frequency of GC B cells (B220^+^IgD^-^GL-7^+^CD95^+^) was expanded in VaxMAP(S) and Alum(S) compared to PBS(S) immunized mice at days 10 and 28 p.i. (**Fig. 3B**). However, at day 60 p.i. GCs were slightly elevated in VaxMAP(S) compared to Alum(S) and PBS(S). These data suggest that VaxMAP(S) drives a prolonged Tfh and GC B cell formation compared to a traditional immunization with Alum(S).

**Fig. 3.**
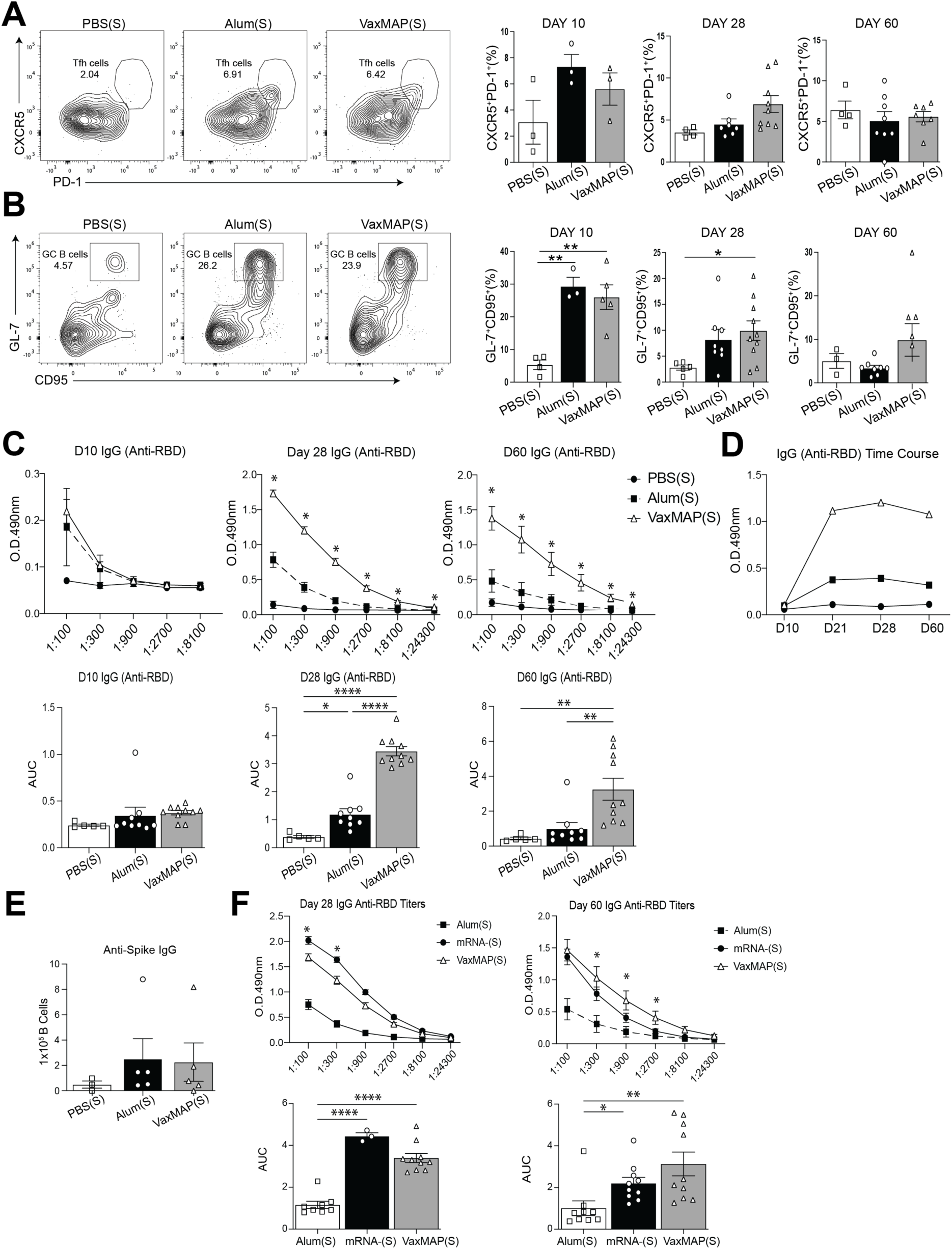
VaxMAP elicits adaptive immune responses. B6 mice were immunized with VaxMAP(S), Alum(S) or PBS(S). (**A**) Tfh cells were stained for flow cytometry and quantified at day 10 post immunization. Representative facs plot (left) and quantification (right) Tfh cells from each group. (**B**) GC B cells were stained for flow cytometry and quantified. Representative facs plot (left) and quantification (right) GC B cells from each group at day 10 post immunization. (**C**) ELISA quantification of serum Anti-RBD IgG antibody of immunized mice at days 10, 28 and 60. Graphs of titers (top) and area under the curve (bottom). (**D**) Anti-RBD IgG antibodies from each group were directly compared over time using a 1:300 titer for each sample from the three time points. (**E**) Representative anti-Spike IgG ELISPOT with number of cells plated per well of B cells from bone marrow of immunized mice. (**F-G**) ELISA quantification of serum Anti-RBD IgG antibody of immunized mice at days 28 or 60 compared to an mRNA vaccine. Data are representative of three independent experiments with 3-8 mice per group. * = p ≤ 0.05, ** = p ≤ 0.01, **** p ≤ 0.0001.

Although VaxMAP(S) and Alum(S) treated mice displayed similar frequency of Tfh and GC B cells at early time points, it may not correlate to the quality of antibody responses. To determine if the “quality” of the humoral response generated by VaxMAP(S) differed amongst vaccination platforms, we assessed the generation of anti-RBD antibodies. As RBD domain of S protein is critical for viral entry, antibodies targeting this domain of SARS-CoV-2 have been shown to contain both neutralizing and protective antibodies (33–36). Sera were collected from mice immunized with VaxMAP(S), Alum(S) or PBS(S) at days 10, 28, and 60 p.i. Akin to the frequency of Tfh and GC B cells, anti-RBD IgG titers from VaxMAP(S) immunized mice were similar to Alum(S) at day 10 but were elevated at day 28 and remained increased compared to both Alum(S) and PBS(S) treated mice at day 60 p.i. (**Fig. 3C**). IgG1 antibodies closely matched total IgG titers, confirming the type 2 biased as previously shown (**Supplemental Fig. 1**) (27). Furthermore, temporal analysis of a single titration from each group demonstrated that while the anti-RBD IgG levels from Alum(S) were higher than PBS(S), VaxMAP(S) had consistently elevated antigen-specific antibodies compared to both groups (**Fig. 3D**). Since antibody secreting B cells in the bone marrow have been shown to be a predictor of long-lasting humoral immunity following immunization (37), we assessed anti-spike secreting B cells in the bone marrow from mice 60 days following immunization by ELISpot (**Fig. 3E**). There was no difference between the numbers of anti-spike secreting B cells from VaxMAP(S) and Alum(S) immunized mice, but both were increased compared to PBS(S). Together these data demonstrate that despite the similar frequency of Tfh and GC B cells, VaxMAP(S) induced elevated and prolonged anti-RBD antibody titers compared to Alum(S).

In current COVID-19 vaccines, antibody titers peak within 2-3 weeks following immunization with a single dose, and then rapidly decline (38). Therefore, multiple boosters have been recommended to patients to prevent reinfection and severe infection during the duration of the pandemic. Given that anti-RBD titers were still elevated 60 days following VaxMAP(S) immunization compared to the other groups, we wanted to compare antibody titers from mice immunized with a lipid-based mRNA vaccine. We obtained sera samples from mice immunized with a single dose of a mRNA-Spike(S) vaccine and compared anti-RBD IgG by ELISA to VaxMAP(S) and Alum(S) at days 28 and 60 post immunization (30). While VaxMAP(S) initially showed a lower humoral response at day 28 compared to mRNA-(S) vaccine, the antibody titers for several dilutions of VaxMAP(S) were higher than mRNA-(S) at 60 days post immunization indicating a more prolonged immune response for VaxMAP(S), which is highly desirable if we want to reduce the number of boosters needed annually (**Fig. 3F**). Thus, VaxMAP(S) enhanced the magnitude and duration of antigen-specific antibody responses when compared to Alum(S) and provided a similar magnitude yet enhanced the duration of antigen-specific antibody responses when compared to mRNA immunizations.

### 3.4. VaxMAP promotes a robust recall response

While the development of a primary adaptive immune response is important following vaccination, a hallmark is to generate a robust early GC and antibody responses upon rechallenge. To determine secondary immune responses mice received a s.c. injection of Alum(S) on into the left flank 65 days after a primary immunization with either VaxMAP(S), Alum(S) or PBS(S) on the right side. Mice were sacrificed 7 days post rechallenge and draining iLNs on the left side were harvested to assess the cellular responses. Tfh cells were similarly expanded in all three groups following rechallenge (**Fig. 4A**). GCs rapidly developed by 7 days following rechallenge, but the percentage of GC B cells was similar in VaxMAP(S) and Alum(S) compared to PBS(S) rechallenged mice, indicating the kinetics in PBS(S) is correlative of a primary response (**Fig. 4B and 3B**). Sera collected from these mice at the time of sacrifice demonstrated elevated anti-RBD-specific IgG levels in VaxMAP(S) and Alum(S) compared to PBS(S) (**Fig. 4C**). Together, these data demonstrate that VaxMAP(S) promotes a robust secondary response upon rechallenge.

**Fig. 4.**
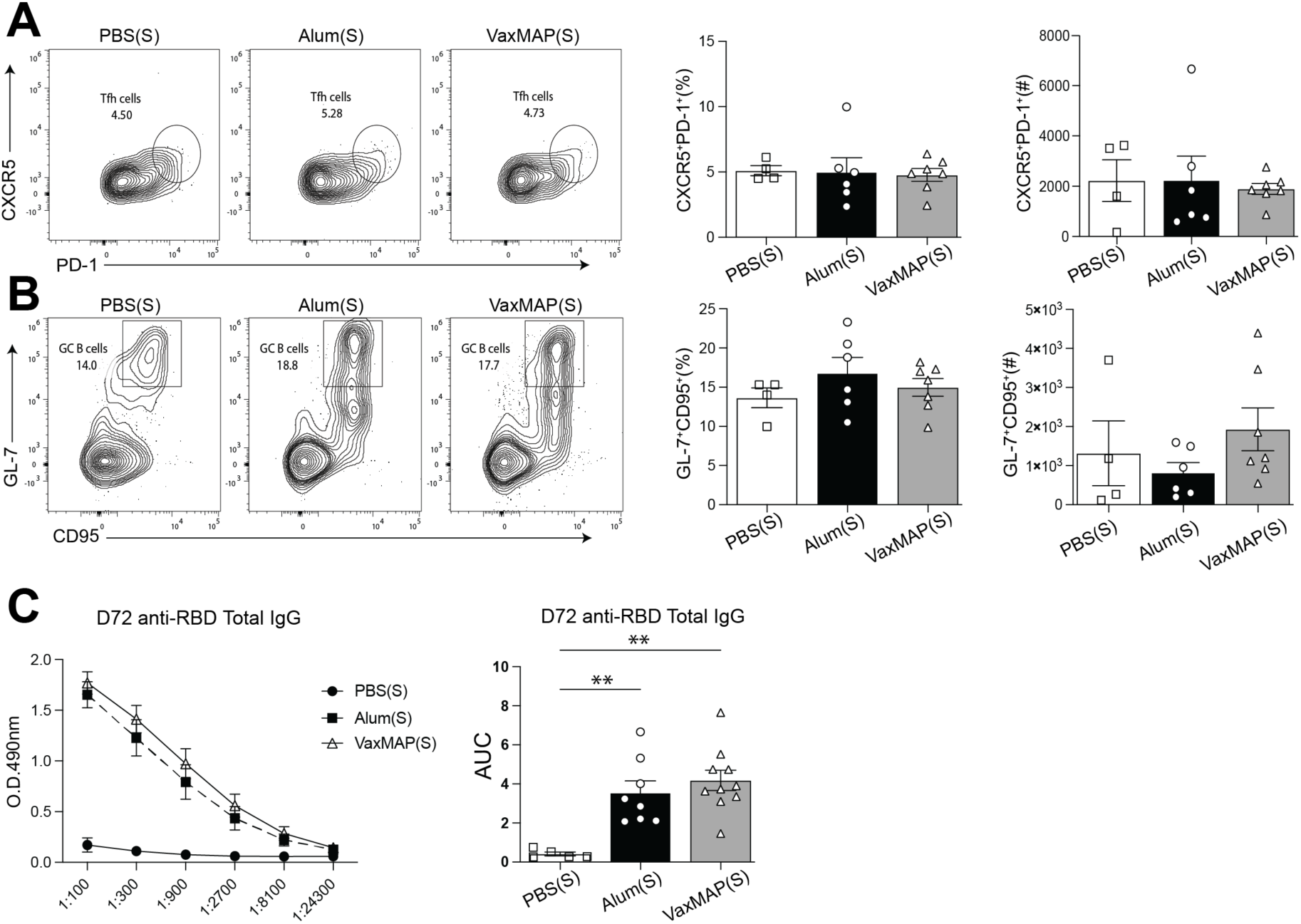
VaxMAP-immunized mice demonstrate a robust recall response. B6 mice were immunized with VaxMAP(S), Alum(S) or PBS(S) and then rechallenged with Alum(S) 65 days later. Draining lymph nodes and sera were collected 7 days after re-immunization. (**A**) Tfh cells were stained for flow cytometry and quantified. Representative facs plot (left) and quantification (right) Tfh cells from each group. (**B**) GC B cells were stained for flow cytometry and quantified. Representative facs plot (left) and quantification (right) GC B cells from each group. (**C**) ELISA quantification of serum Anti-RBD IgG antibody of re-immunized mice. Graphs of titers (left) and area under the curve (right). Data are representative of three independent experiments with 5-8 mice per group. ** = p ≤ 0.01.

### 3.5. VaxMAP elicits protection against influenza infection

While antigen loaded VaxMAP could drive primary and secondary immune responses, we next wanted to assess whether it can elicit protection against a live infection following immunization. Given that SARS-CoV-2 requires BSL3 containment, and the mouse strains required for infections are not comparable to humans, we chose to test VaxMAP against a murine influenza A virus infection (H1N1 A/PR8/34 strain (PR8)) (39, 40). We first wanted to validate MAP as a vaccine depot using recombinant PR8 antigen, so we added viral glycoprotein hemagglutinin (HA) to VaxMAP as it is also the most common target of influenza neutralizing antibodies(41). HA is a type I fusion protein similar to spike and mediates receptor binding and virus fusion. Therefore, we loaded HA antigen into MAP following the same strategy as for VaxMAP with spike. We immunized mice with either VaxMAP(HA), Alum mixed with HA (Alum(HA)), PBS (HA) or MAP alone (no antigen). We collected sera from the mice at days 21, 45 and 55 p.i. and assessed anti-HA IgG antibodies from the immunized mice. VaxMAP(HA) and Alum(HA) had similar anti-HA IgG titers through 45 days, but the antibody levels in VaxMAP(HA) remained elevated at day 55 p.i. (**Fig. 5A**). To determine the quality of the antibodies generated from each group we performed an HA-inhibition assay. A lower titer of sera from VaxMAP(HA) mice was required to inhibit HA agglutination of red blood cells compared to Alum(HA) immunized mice, indicating these antibodies were better at neutralizing HA (**Fig. 5B**). Next, to assess protection against a live virus we administered a high dose of influenza to the mice immunized 60 days earlier. Mice immunized with VaxMAP(HA) showed reduced weight loss and began to rebound faster compared to Alum(HA)-treated mice, and both experienced decreased weight loss compared to the PBS (HA) and MAP injected mice (**Fig. 5C**). VaxMAP(HA) immunized mice displayed better survival against a challenge with a high dose of influenza compared to all other groups (**Fig. 5D**), suggesting VaxMAP(HA) elicited better protection than Alum(HA). Overall, these results demonstrate that VaxMAP loaded with an influenza-specific antigen is capable of providing better neutralizing antibodies resulting in increased protection against infection compared to Alum, an adjuvant commonly used in influenza vaccines. Furthermore, the fact that the humoral response with VaxMAP(HA) aligns with that observed with VaxMAP(S) strongly suggests that VaxMAP(S) would also be effective against a SARS-CoV-2 infection.

**Fig. 5.**
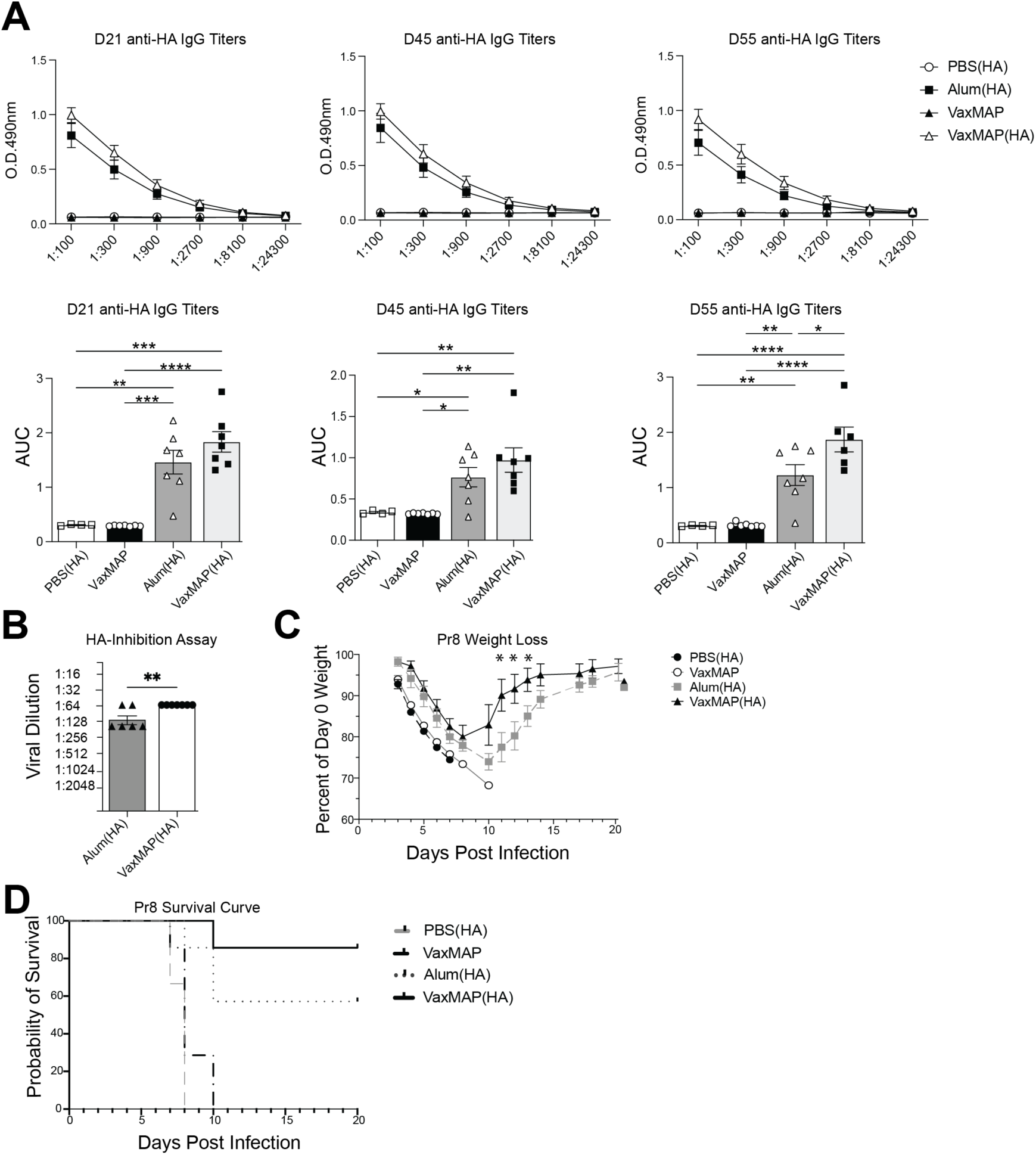
VaxMAP elicits protection against influenza infection. **(A)** B6 mice were immunized with VaxMAP(HA), Alum(HA), PBS(HA), or empty VaxMAP and ELISA quantification of serum Anti-RBD IgG antibody of immunized mice at days 21, 45 and 55. Graphs of titers (top) and area under the curve (bottom). (**B**) Sera from day 55 immunized mice was used in a HA agglutination assay with different dilutions of PR8. (**C-D**) Mice were given a high dose of PR8 60 days after immunization (**C**) Daily weight loss based on the percentage of the initial weight of each mouse out to 20 days post infection. (**D**) Survival curve of mice following infection. Data are representative of 2 independent experiments with 5-8 mice per group. * = p ≤ 0.05, ** = p ≤ 0.01, *** = p ≤ 0.001, **** p ≤ 0.0001.

## 4. CONCLUSION

In summary, this work has established the ability of VaxMAP to elicit a strong immune response and protect against a viral infection. Furthermore, VaxMAP has the potential as a platform technology that can be readily deployed for various pathogens and potentially be used as a depot to deliver multiple antigens simultaneously. Based on the recent pandemic and the almost certain emergence of other novel pathogens, the ability to rapidly generate effective immunity in patients as pathogens emerge will be critical to quickly develop population level immunity which can help alleviate ripple effects like the current economic crisis. Taken together, VaxMAP demonstrated the ability to enhance the adjuvant effects of a biomaterial platform to generate more robust and maintained antibody responses and improved protection against a viral infection when compared to traditional platforms. Currently, VaxMAPs provides a polarized type 2 immune response generating a strong IgG1 response, which is important in for the neutralization of toxins and impairing viral attachment. Thus, it would be important next to demonstrate that the ability to tune VaxMAP to impact type 1 immune responses by adding in TLR7 agonists or alter the biomaterial to drive this response, strengthening the role for using modular injectable biomaterials as a vaccine application.

## ACKNOWLEDMENTS

Research reported in this publication was supported by the National Science Foundation (NSF) under award number 2031727. J.S.W and laboratory were also supported in-part by the National Institutes of Health (NIH) R01 AR073912. We would like to acknowledge S. Eisenbarth at Yale University School of Medicine for providing serum samples from mice immunized with mRNA-1273 vaccine. J.S.C. was supported by NIH grants T32GM136651 and F30HL149151.

## AUTHOR CONTRIBUTIONS

D.P.M. designed and performed experiments and wrote the manuscript; J.S.W. designed and performed experiments and wrote the manuscript. S.D. designed and performed experiments and wrote the manuscript. M.N., D.A., and J.S.C. performed experiments. W.M.W and P.O.S designed experiments and wrote the manuscript. All authors read and approved the manuscript.

---

**Table S1.**
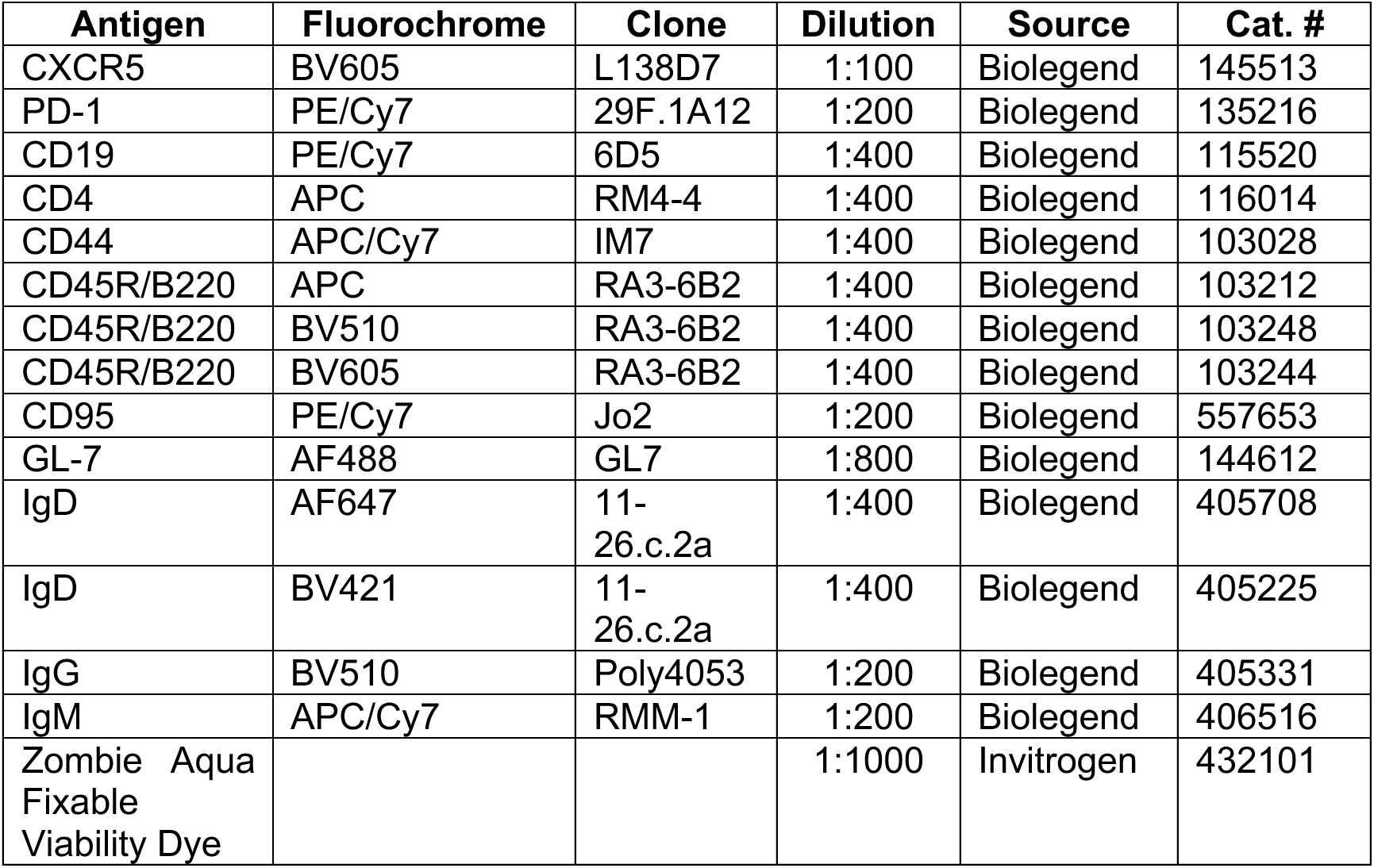
Antibodies used for flow cytometric assays.

