## Supplemental Figure 1 for "A novel microporous biomaterial vaccine platform for long-lasting antibody mediated immunity against viral infection"

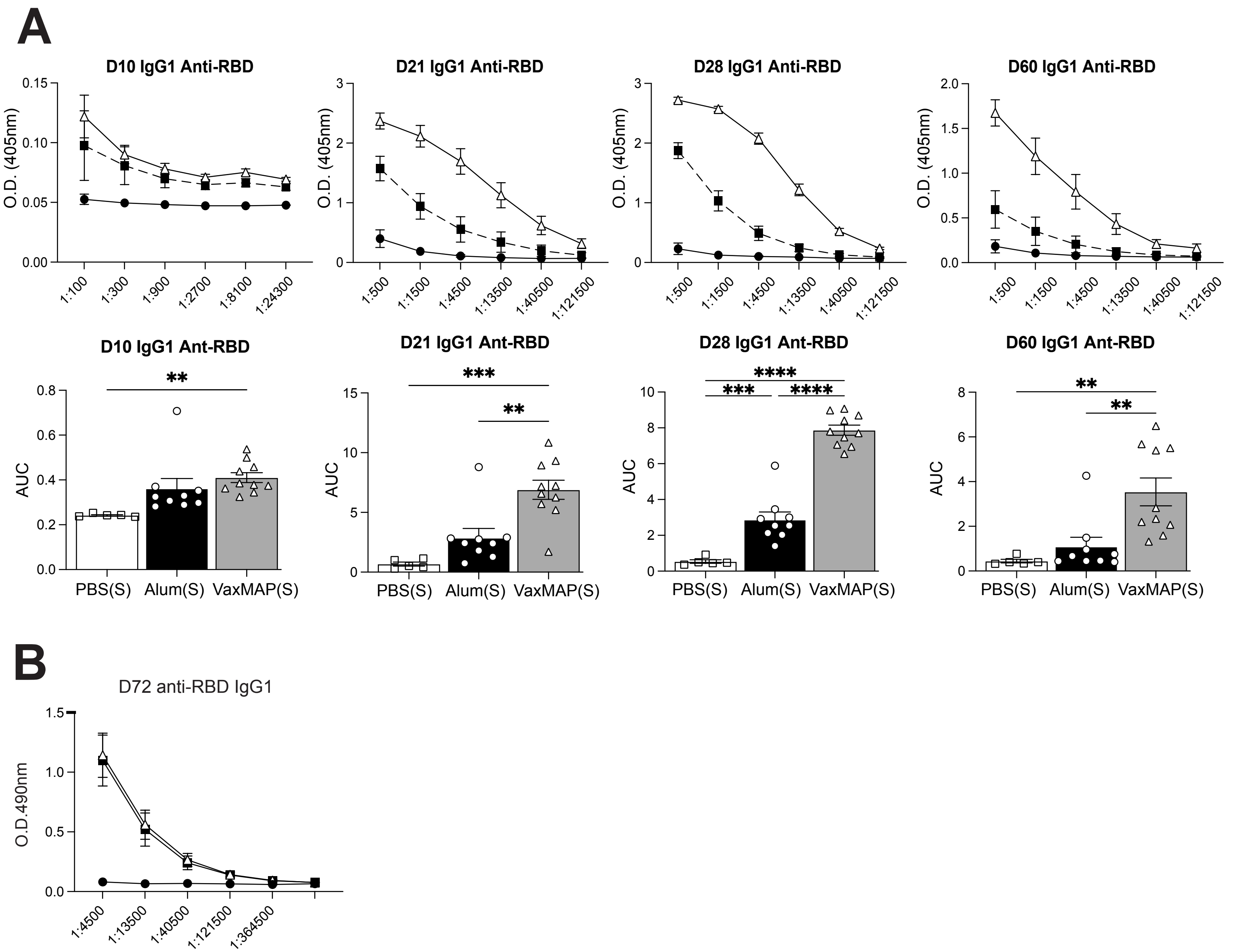


**Supplemental Figure 1 VaxMAP elicits a strong IgG1 antigen-specific antibody response.** B6 mice were immunized with VaxMAP(S), Alum(S) or PBS(S). To determine the type 2 immune response driven by Alum and VaxMAP, (**A**) ELISA were perfromed to quantify the serum Anti-RBD IgG1 antibody of immunized mice at days 10, 28 and 60. Graphs of titers (top) and area under the curve (bottom). (**B**) B6 mice were immunized with VaxMAP(S), Alum(S) or PBS(S) and then rechallenged with Alum(S) 65 days later. ELISA were perfromed to quantify the serum Anti-RBD IgG1 antibody of immunized mice 7 days post rechallenge. Data are representative of three independent experiments with 3-8 mice per group. * = p ≤ 0.05, ** = p ≤ 0.01, **** p ≤ 0.0001.
